# SUMO modulates meiotic crossover rates between and within vertebrate species

**DOI:** 10.64898/2026.03.26.714374

**Authors:** S. Lava Kumar, Rohit Beniwal, Aradhana Mohanty, Ajith Kumar, Anjali Kumari, R. K. Gandham, Neil Hunter, H.B.D. Prasada Rao

**Affiliations:** BRIC-National Institute of Animal Biotechnology, Hyderabad, Telangana 500032, India; Graduate studies, Regional Center for Biotechnology, Faridabad - 121 001, India; Howard Hughes Medical Institute, University of California Davis, Davis, CA 95616, USA; Department of Microbiology & Molecular Genetics, University of California Davis, Davis, CA 95616, USA

## Abstract

Crossing over during meiosis drives genetic diversity and ensures the accurate segregation of homologous chromosomes. Variation in the rate of crossing over has been linked to evolutionary divergence and environmental adaptability, shaping fitness and responses to selective pressures. Despite its significance, the molecular mechanisms underlying this variation remain poorly understood. Crossover sites are selected from a large pool of potential sites initiated by programmed DNA double-strand breaks. Post-translational modification by SUMO (Small Ubiquitin-like Modifier) has been implicated in this process. Here, we show that crossover rate, chromosome length, and abundance of chromosome-associated SUMO are positively correlated across a range of vertebrate species, including mouse, chicken, pig, cattle, sheep, and goat. Crossover variation between goat breeds across the Indian subcontinent was also positively correlated with chromosomal SUMO level. Furthermore, modulating SUMO levels in cultured goat spermatocytes altered crossover frequency. Cumulatively, these observations point to a central role for SUMO in mediating crossover variation both between and within vertebrate species.

## Introduction

Meiosis produces haploid gametes during sexual reproduction ^1^. In meiotic prophase I, crossing over between homologous chromosomes promotes accurate segregation and genetic variation^2^. Crossover rates vary between individuals, sexes, and species^3^. A four-locus model suggests that crossover variation is beneficial for environmental adaptability^4^. Notably, a population with a higher recombination rate shows a more robust response to natural and artificial selection pressures^5–7^. This is manifest in domesticated plant and livestock species, which have higher crossover rates than their predecessors or wild counterparts^8,9^. Although crossover rates vary between individuals and species, the distribution of crossovers is controlled to ensure that at least one crossover forms between each chromosome pair^10^. These obligate crossovers are required to ensure accurate disjunction of homologous chromosomes at the first meiotic division^10^.

Crossing over is one outcome of homologous recombination, which is initiated during meiosis by SPO11-catalyzed DNA double-strand breaks (DSBs)^11^. DSB ends are resected to form long 3’ single-stranded tails that assemble filaments of the RAD51 and DMC1 recombinases to catalyze homology search and DNA strand invasion^2,12^. Nascent strand invasion intermediates are stabilized by a conserved set of meiosis-specific ZMM factors that facilitate chromosome pairing and synapsis and enable the differentiation of crossover sites. Initially defined in budding yeast, the ZMMs include *Zip1^Sycp1^*, *Zip2^Shoc1^*, *Spo16*, *Zip3^Rnf212^*, *Zip4^Tex11^*, *Mer3^Hfm1^*, and *Msh4*-*Msh5* (names of mammalian counterparts are superscript)^2,13,14^. Among these, the RING-family E3 ligase RNF212 plays a key role in crossover designation by localizing along the synaptonemal complex (SC) and becoming selectively enriched at a subset of recombination sites, where it stabilizes recombination intermediates. In parallel, post-translational modification by SUMO (Small Ubiquitin-like Modifier) acts in concert with RNF212 to regulate recombination progression. SUMO localizes as discrete foci along chromosome axes during early prophase I, particularly during zygonema, in association with the SC, and this axis-associated SUMO is largely dependent on RNF212. Together, RNF212 and SUMO are thought to stabilize recombination intermediates and promote their differentiation into crossovers^15,16^. At designated crossover sites, ZMMs and crossover-specific factors such as MutLγ (MLH1-MLH3) facilitate the formation and resolution of double Holliday junction (dHJ) intermediates specifically into crossovers^17,18^. At other recombination sites, the invading strand is extended by DNA synthesis before being dissociated and annealing to the other DSB end to produce a non-crossover, without exchange of chromosome arms^2,18^. ^16^Genetic, epigenetic, and environmental variation can modulate the rate of crossing over^19–21^. The rapid evolution of meiotic crossover landscapes is exemplified by the lack of conservation between human and chimpanzee recombination maps^22^. Furthermore, crossover rate is not dictated solely by karyotype or genome size: for example, goats and cattle have the same number of chromosome arms and similar genome sizes but different recombination rates^23,24^. Also, sex-specific variation in crossover rate occurs in many species, often with females having a higher rate than males, although the opposite is also common, as seen in sheep, cows and deer^25–27^. Genome-wide association studies (GWAS) in humans, mice, cattle, pigs, sheep, and goats have linked several genetic variants to heritable differences in recombination rates^28–32^. In mice and humans, the numbers of DSBs and crossovers are positively correlated with the axial lengths of meiotic prophase chromosomes, measured as the lengths of synaptonemal complexes^12,20,33,34^. However, DSB frequency is not the sole determinant of the crossover rate. For example, mice with reduced DSB levels (via reduced SPO11 activity) can maintain a wild-type crossover rate through a process termed crossover homeostasis^35^. In mammals, a minor but variable fraction of DSBs is matured into crossovers. For example, in mouse spermatocytes, only ∼10% of DSBs mature into crossovers, whereas in humans, ∼35% of DSBs become crossovers^20,27,36^. How such variation in crossover bias is acquired during evolution remains unclear. In this study, using a variety of vertebrate species, we provide evidence for the important role of SUMO in crossover variation.

## Results

### Crossover rate and chromosome-axis length are correlated across species defining a quasi-fixed unit of axis length per crossover

To understand what drives variation in crossover rates among vertebrates, we examined how crossover frequency relates to genome size, karyotype, and chromosome axis length during meiotic prophase I in evolutionarily distinct domesticated livestock species: chicken (*Gallus gallus domesticus*), pig (*Sus scrofa*), cattle (*Bos indicus*), sheep (*Ovis aries*), and goat (*Capra hircus*), along with mouse (*Mus musculus*) as a well-established model (Fig. 1A). Crossovers were quantified by immunostaining surface-spread spermatocyte nuclei for the crossover marker MLH1 and the chromosome axis protein SYCP3 (Fig. 1B). MLH1 foci were quantified in late-pachytene stage nuclei (Fig. 1C, Fig. S1B), providing a reliable cytological estimate of crossover numbers. Consistent with previous studies, MLH1 counts closely matched known genetic map lengths across species, validating our approach.^37,38,38^

**Fig. 1.**
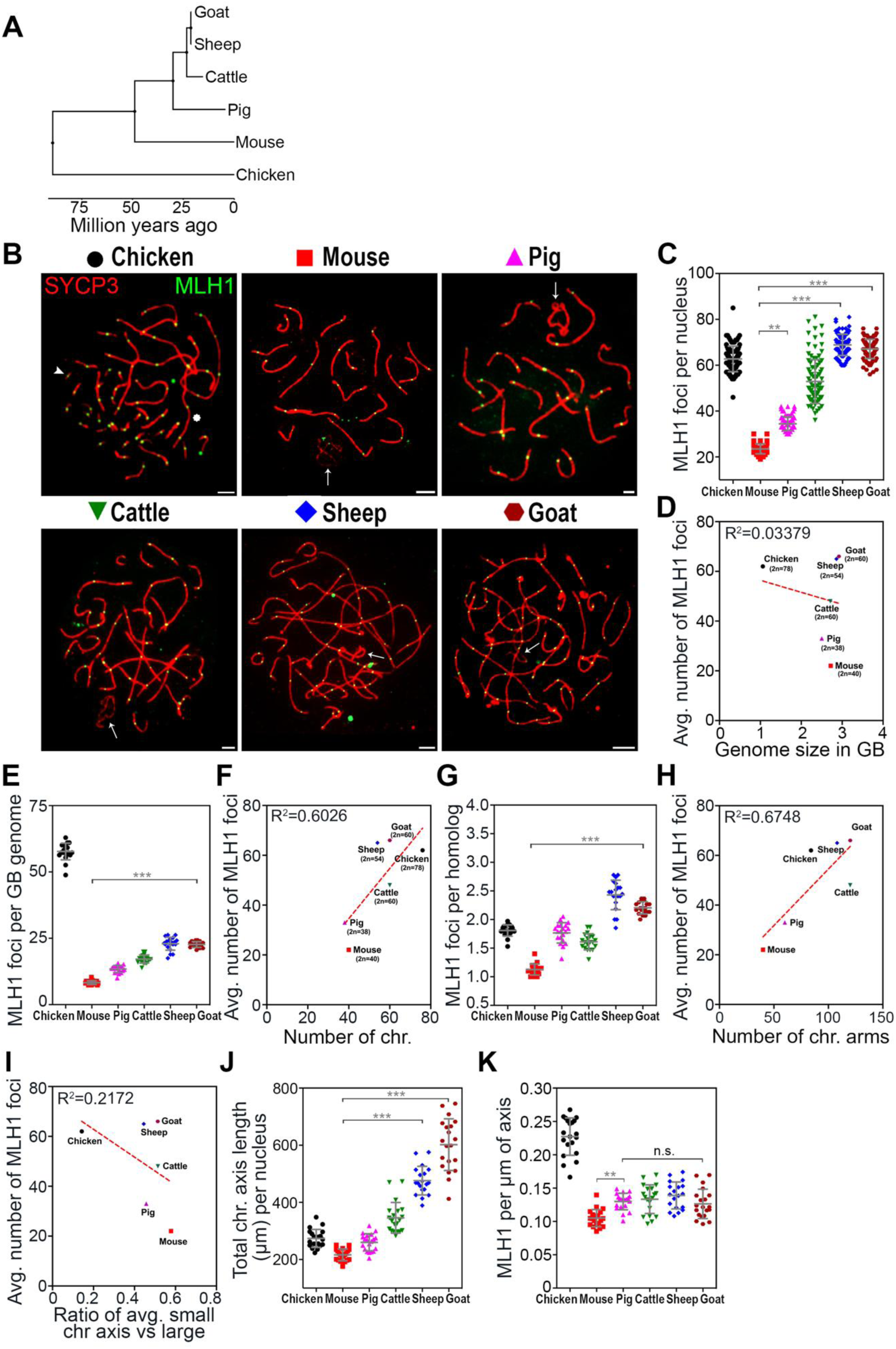
Comparative analysis of MLH1-defined crossovers across species. (A) A cladogram based on meiotic recombination genes illustrating evolutionary relationships among the analyzed species, with chicken as the outgroup followed by mouse, pig, cattle, sheep, and goat. (B) Representative late pachytene spermatocyte nuclei from each species, immunostained for the chromosome axis protein SYCP3 (red) and the crossover marker MLH1 (green). (C) Quantification of MLH1 foci per nucleus across species. (D) Relationship between the average (Avg.) number of MLH1 foci and genome size (see Table 1). (E) MLH1 foci normalized to genome size, expressed as foci per gigabase (Gb). (F) Correlation between the average number of MLH1 foci and chromosome number (Chr.) (see Table 1). (G) MLH1 foci normalized per homologous chromosome pair. (H) Relationship between the average number of MLH1 foci and chromosome arm number (see Table 1). Chromosome arms were defined based on centromere position. (I) Correlation between MLH1 foci and the ratio of small to large chromosome axes in each species. Chromosome size classes were defined as follows: chicken (29 small, 10 large pairs), mouse (4 small, 16 large pairs), pig (6 small, 13 large pairs), cattle (14 small, 16 large pairs), sheep (13 small, 14 large pairs), and goat (17 small, 13 large pairs). (J) Total synaptonemal complex (chromosome axis) length per nucleus. (K) MLH1 foci density along chromosome axes, expressed as foci per micron of synaptonemal complex length. Error bars represent mean ± s.d. Statistical significance was assessed using a two-tailed Mann–Whitney test (***P ≤ 0.001, **P ≤ 0.01, n.s., not significant). Sex chromosomes are indicated by arrows. In chicken, arrowheads denote microchromosomes and asterisks indicate the largest chromosome. Coefficients of determination (R²) and linear regression (red dashed lines) are shown where applicable. Scale bars, 10 μm.

**Table 1:** Cross-species comparison of recombination markers and chromosome features:

| Species | Chr.no.<br>(2n) | Centromere<br>location | Genome<br>Size<br>(GB) | MLH1<br>foci<br>(avg.) | Rad51<br>foci<br>(avg.) | RPA<br>foci<br>(avg.) | RNF212<br>foci<br>(avg.) | Avg.<br>chromosome<br>length<br>(µm) | Avg.<br>MLH1<br>per<br>µm | Chr. Arms<br>(calculated<br>based on<br>centromere<br>location) |
| --- | --- | --- | --- | --- | --- | --- | --- | --- | --- | --- |
| Chicken | 78 | 68- Telocentric<br>4-Submetacentric<br>2- Acrocentric<br>2- Metacentric | 1.06 | 62 | 225 | 404 | 140 | 273 | 0.227 | 84 |
| Mouse | 40 | 40- Telocentric | 2.72 | 23 | 220 | 218 | 112 | 216 | 0.105 | 40 |
| Pig | 38 | 10-Submetacentric<br>4-Subtelocentric<br>12- Metacentric<br>12- Telocentric | 2.50 | 34 | 83 | 153 | 143 | 259 | 0.130 | 60 |
| Cattle | 60 | 58- Acrocentric<br>1-Submetacentric<br>1- Acrocentric | 2.82 | 52 | 137 | 242 | 110 | 348 | 0.133 | 120 |
| Sheep | 54 | 6-Submetacentric<br>46- Acrocentric<br>1- Acrocentric<br>1-Submetacentric | 2.86 | 68 | 187 | 259 | 293 | 475 | 0.139 | 108 |
| Goat | 60 | 58- Acrocentric<br>1- Acrocentric<br>1-Submetacentric | 2.92 | 67 | 186 | 308 | 270 | 618 | 0.126 | 120 |

For example, chicken spermatocytes exhibited 62 ± 5.5 MLH1 foci per nucleus, corresponding to a genetic map length of ∼3150 cM, whereas mice showed 23 ± 2.1 foci, consistent with ∼1150 cM^39,40^. Similar agreement was observed across other species: in pig, 34 ± 3.0 MLH1 foci matched a genetic map length of ∼1747 cM (∼35 expected); in cattle, sex-averaged *Bos indicus* maps (∼3160 cM) corresponded to 52 ± 9.5 (∼39 MLH1 foci expected in *Bos taurus* with genetic map length ∼1975 cM), respectively; in sheep (*Ovis aries*), 68 ± 4.6 MLH1 foci aligned with a genetic map length of ∼3817 cM (∼76 expected); and in goat (*Capra hircus*), 67 ± 4.1 MLH1 foci matched ∼3500 cM (∼70 expected)^41–45^.

Possible drivers of the different crossover rates in vertebrate species include differences in genome size (Mb of DNA), the number of chromosomes and chromosome arms, karyotype asymmetry (the size difference between the largest and smallest chromosomes), and the physical lengths of chromosomes during meiotic prophase I. To evaluate these possibilities, we systematically compared each of these parameters with crossover frequency across the species analyzed.

#### Genome size

Total genome size was not well correlated with the number of MLH1 foci per nucleus (Fig. 1D; *R*^2^ = 0.03379). For instance, chicken, with a relatively small genome (∼1.2 Gb), exhibited a similar number of MLH1 foci as sheep and goat, which have much larger genomes (∼2.6–2.7 Gb). At the same time, chicken displayed substantially higher crossover numbers than mouse, pig, and cattle, despite these species having comparable genome sizes (∼2.5–2.7 Gb). When normalized to genome size, the number of MLH1 foci per Gb of DNA further highlighted this lack of scaling. Chicken showed a markedly elevated crossover density, with ∼2.5–7-fold higher MLH1 foci per Gb compared to other species (Fig. 1E). In contrast, the remaining species displayed more moderate variation: pig (∼1.6–4-fold), cattle (∼2–3.3-fold), sheep (∼2.7-fold), and goat (∼2.5-fold differences relative to each other), while mouse consistently showed lower values. Overall, MLH1 density ranged from ∼0.36 to 2.73 foci per Gb across species, indicating that genome size alone is not a major determinant of crossover rate.

#### Chromosome number, structure, and asymmetry

Crossover assurance ensures that each pair of homologs becomes connected by at least one obligatory chiasmata, as required for accurate homolog segregation at the first meiotic division. Thus, chromosome number is expected to be one driver of crossover rate. Indeed, MLH1 foci per nucleus and chromosome number were positively correlated across species (Fig. 1F; *R^2^* = 0.6026; Table 1). However, the fit was not good and species with similar numbers of chromosomes, such as mouse and pig (20 and 19, respectively) or goat and cattle (30), had significantly different numbers of MLH1 foci per nucleus (Fig. 1F) and per chromosome (Fig. 1G; Table 1), indicating additional drivers of crossover rate beyond chromosome number. Differences in the number of chromosome arms between species did also not account for these deviations (Fig. 1H; *R^2^* = 0.6748). Mechanisms that help ensure crossing over occurs between the smallest chromosomes might also drive a higher overall crossover rate. Therefore, we asked whether karyotype asymmetry (specifically the ratio of the largest chromosome to the smallest chromosome) was correlated with the number of MLH1 foci, but the fit was poor (Fig. 1I; *R^2^* = 0.2172).

#### Chromosome axis length

Covariation of crossover rate and prophase I chromosome-axis length was previously reported in mice and humans^46^. Therefore, we compared the axis/synaptonemal complex (SC) lengths in late-pachytene spermatocytes from mouse, chicken, pig, cattle, sheep and goat. Chromosome axis lengths were strongly positively correlated with the numbers of MLH1 foci per nucleus (Fig. 1J, *R^2^* = 0.8984), with chicken being the sole outlier. Moreover, plotting the frequency of MLH1 foci per µm of SC suggests a quasi-fixed crossover rate per unit-length of 0.10–0.22 MLH1 foci per µm of axis (Fig. 1K).

A clear exception to this trend is chicken, with 0.22 MLH1 foci per µm SC, indicating an additional driver of crossover rate in this species. A likely candidate is the obligate crossovers between the numerous very small chromosomes in chicken, with 29 out of 39 chromosomes being classified as micro- or dot-chromosomes^47^. Consistently, if the micro- and dot-chromosomes are excluded from our analysis, the ratio of MLH1 per µm axis is now more similar to that of other vertebrate species (0.16 ± 0.02 MLH1 per µm SC; Fig. S1A 1K).

### The number of recombination precursors per crossover is similar across livestock species

In most species, a minor fraction of precursor DSBs mature into crossovers^2^. To estimate DSB numbers and understand the relationships with crossover frequency and chromosome axis length, spermatocyte nuclei were immunostained for the DNA strand-exchange protein RAD51, to mark DSB sites, and SYCP3 to mark chromosome axes and stage nuclei (Fig. 2A). Although the numbers and dynamics of RAD51 foci varied across species (Fig. S2A and B), the ratio of RAD51 foci per MLH1 focus (≈DSBs per crossover) was similar in all livestock species, averaging 2.8 (range 2.486 to 2.977, Fig. 2B), i.e. ∼35% of DSBs are converted into crossovers, which is similar to human spermatocytes^48^. The outlier was mouse, with ∼10 RAD51 foci per MLH1, confirming that the crossover/noncrossover ratio is not a fixed parameter in mammalian meiosis. The density of RAD51 per µm of SC was also similar for pig, cattle, sheep, and goat at ∼0.354 (Fig. 2C), but was significantly higher for chicken (0.823) and mouse (1.053). Thus, DSB density does not appear to be a fixed function of axis architecture. These inferences were confirmed by staining for a second DSB marker, the single-stranded DNA binding protein, RPA^15^ (Fig. 2D–F and S3A, B).

**Fig. 2.**
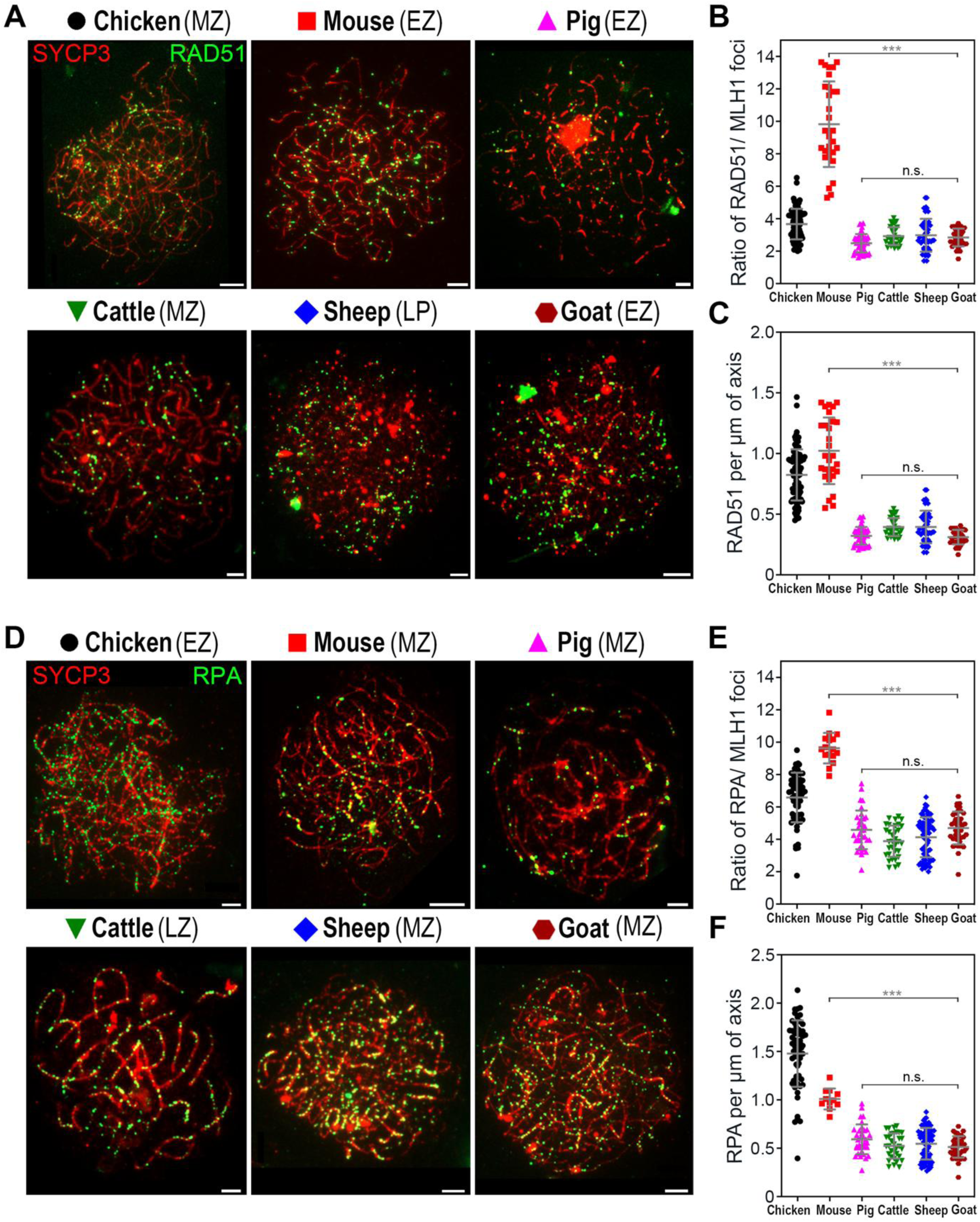
RAD51 and RPA dynamics reveal conserved DSB-to-crossover ratios. (**A**) Spermatocyte nuclei from indicated species immunostained for SYCP3 (red) and RAD51 (green). (**B**) Quantification of ratio of RAD51 per MLH1 foci. (**C**) RAD51 foci density along chromosome axes, expressed as foci per micron of synaptonemal complex length. (**D**) Spermatocyte nuclei from indicated species immunostained for SYCP3 (red) and RPA (green). (**E**) Quantification of ratio of RPA per MLH1 foci. (**F**) RPA foci density along chromosome axes, expressed as foci per micron of synaptonemal complex length. Error bars show mean ± s.d. (***) *P* ≤ 0.001, (n.s.) *P* ≥ 0.1, two-tailed Mann-Whitney. LP-Leptonema, EZ- Early zygonema, MZ- Mid zygonema, LZ-Late zygonema. Scale bars = 10μm.

### RNF212 foci positively correlates with axis length and crossover rate

In mouse, RNF212 is required for the differentiation of crossover sites and the chromosomal localization of crossover-specific factors such as MutLγ^16,49^. RNF212 localizes specifically to the SC central region, initially as numerous foci before accumulating at crossover sites during mid-late pachytene while diminishing elsewhere^16^. Immunostaining of RNF212 revealed that this dynamic behavior of RNF212 is conserved across vertebrates (Fig. 3A and B). Also, the density of RNF212 foci in early pachytene spermatocytes was similar across the species tested, at 0.4–0.6 foci per µm of SC (Fig. 3C), possibly reflecting a conserved structural feature of the SC central region. Consequently, numbers of RNF212 foci were positively correlated with both SC/axis length (Fig. 3D) and the numbers of MLH1 foci across vertebrate species (Fig. 3E). The number of RNF212 foci per crossover was similar for mouse, pig, sheep and goat (∼4 RNF212 per MLH1) but lower for chicken and cattle (∼2 foci) (Fig. 3F). RNF212 foci did not no correlated with early precursors (RAD51 and RPA) (Fig. 3G and H).

**Fig. 3.**
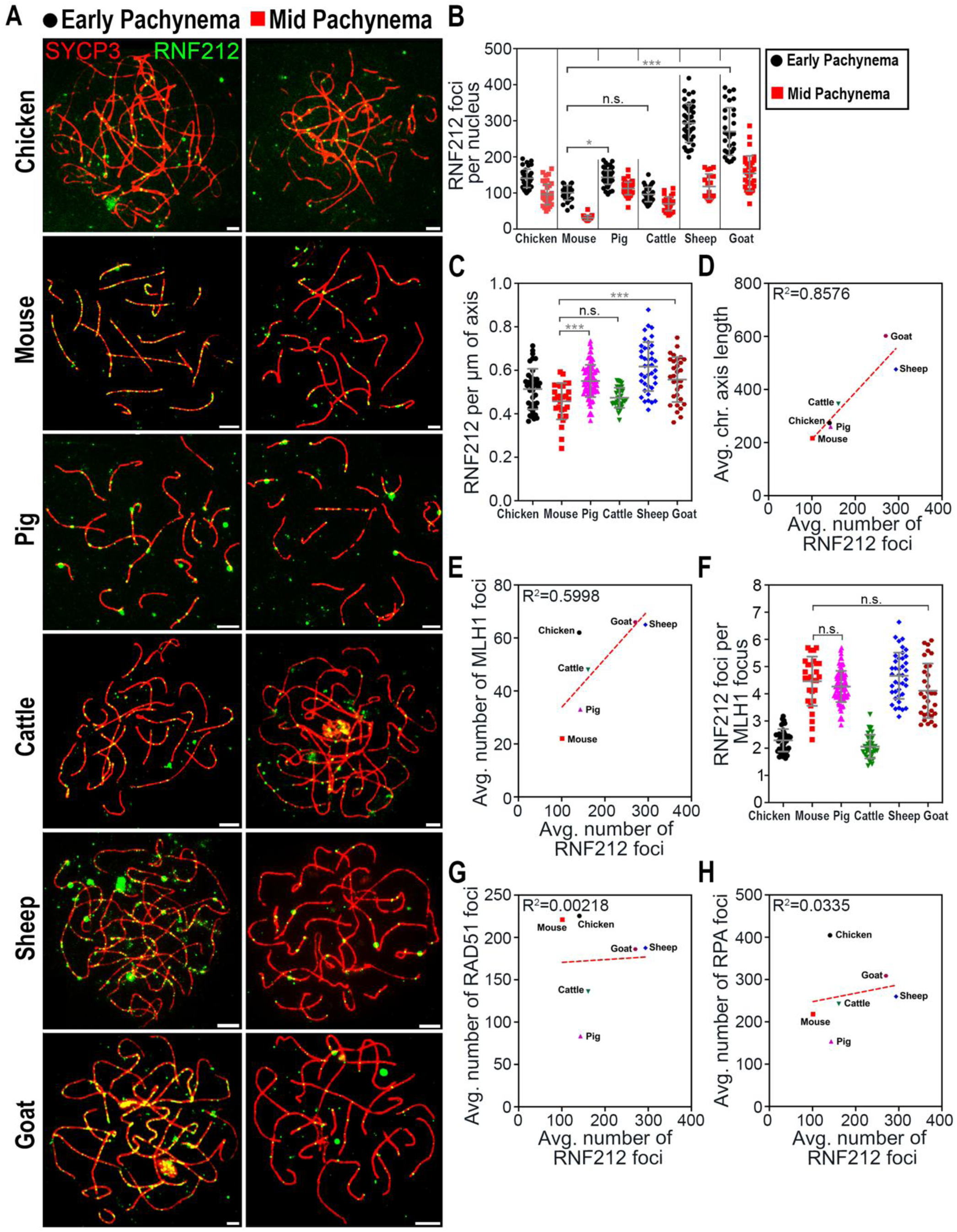
Cross-species analysis of RNF212 and crossover designation. (**A**) Representative spermatocyte nuclei from the indicated species immunostained for the chromosome axis protein SYCP3 (red) and the crossover designation factor RNF212 (green) across the indicated meiotic substages. (**B**) Quantification of RNF212 foci in respective stages. (**C**) RNF212 foci density along chromosome axes, expressed as foci per micron of synaptonemal complex length. (**D**) Comparison of average chromosome axis length with average number of RNF212 foci. (**E**) Comparison of the average number of MLH1 with the average number of RNF212 foci. (**F**) Quantification of RNF212 foci per MLH1 focus. (**G**) Comparison of the average number of RAD51 foci with the average number of RNF212 foci. (**H**) Comparison of the average number of RPA foci with the average number of RNF212 foci. Error bars show mean ± s.d. (***) *P* ≤ 0.001, (*) *P* ≤ 0.1, (n.s.) *P* ≥ 0.1, two-tailed Mann-Whitney. R^2^ correlation values are included in graphs along with red dashed line wherever analysed. Scale bars = 10μm.

### Axis-associated SUMO positively correlates with SC length and crossover rate

Previous studies in mouse showed that SUMO localizes as a focal pattern along chromosome axes as synapsis ensues during zygotene, before disappearing during pachytene while accumulating at centromeric heterochromatin and the XY chromosomes (to mark the transcriptionally silenced sex body) (Fig. 4A)^15^. Moreover, axis-associated SUMO was show to be largely RNF212 dependent^15^. The patterns and dynamics of SUMO1 immunostaining in spermatocyte nuclei from livestock species were similar but also distinct from that of mouse (Fig. 4A and B). Most notably, unlike in mouse, axis-associated foci of SUMO1 persisted during pachytene, albeit at lower levels than seen in zygotene. The numbers of SUMO1 foci were also positively correlated with both axis length and numbers of MLH1 foci (Fig. 4C and 4D). Also, the density of SUMO1 foci in late zygotene spermatocytes was similar across pig, cattle, sheep and goat, at 0.5–0.8 foci per µm of SC (Fig. 4E), lower in chicken (0.4 foci) and higher in mouse (1.1 foci). Similarly, the number of SUMO1 foci per MLH1 focus was also similar in pig, cattle, sheep and goat (∼5-6) across species, lower in chicken (∼2) and higher in mouse (∼11) (Fig. 4F). Overall, these results suggest that axis-associated SUMO positively correlates with crossovers and chromosome length in higher-order vertebrates.

**Fig. 4.**
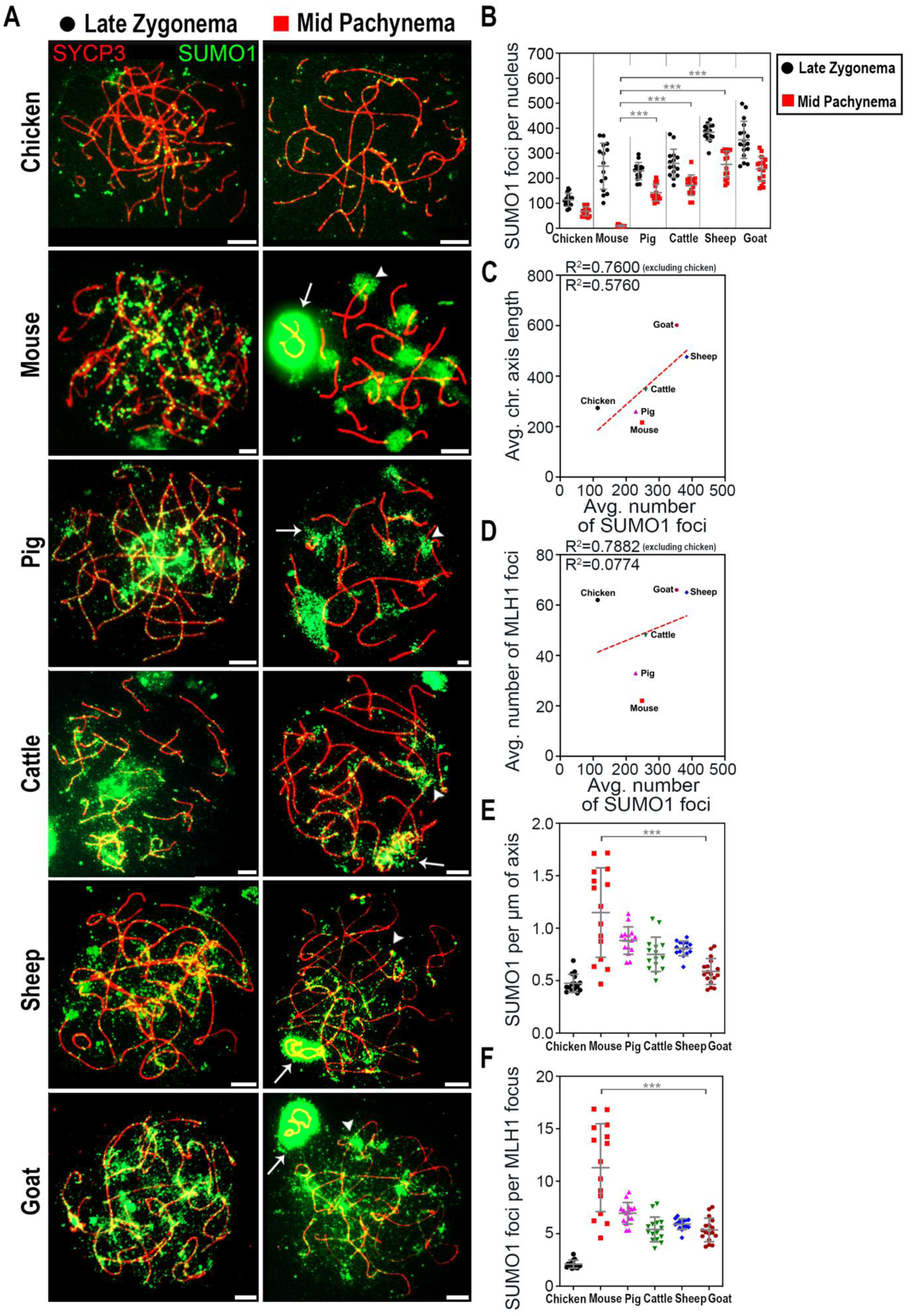
Cross-species analysis of SUMO1 and its relationship with crossovers. (**A**) Spermatocyte nuclei from indicated species immunostained for SYCP3 (red) and SUMO1 (green) in all respective stages. (**B**) Quantification of immunostaining foci in respective stages. (**C**) Comparison of average chromosome axis length with average number of SUMO1 foci. (**D**) Comparison of average number of MLH1 foci with average number of SUMO1 foci. (**E**) Quantification of SUMO1 per micron of chromosome axis in late zygonema nuclei. (**F**) Quantification of SUMO1 foci per MLH1 focus in late zygonema nuclei. Error bars show mean ± s.d. (***) *P* ≤ 0.001, two-tailed Mann-Whitney. R^2^ correlation values are included in graphs along with red dashed line wherever analysed and shown by excluding chicken. Scale bars = 10μm.

### Intra-species variation in crossover rate positively correlates with chromosome axis length and levels of axis-associated SUMO in goat breeds

The goat breed used in the analysis thus far is Jamunapari. However, 43 domestic goat breeds are recognized on the Indian subcontinent, (National Bureau of Animal Genetic Resources-India) that differ in their fitness and reproductive characteristics. We collected spermatocytes from five of these breeds and immunostained them for MLH1 and SYCP3 (Fig.5A). MLH1 focus numbers varied significantly, with averages of 51 ± 6.7 SD in Osmanabadi; 61 ± 3.4 SD in Jamunapari; 66 ± 3.2 SD in Nandidurga; 70.1 ± 5.1 SD in Mahbubnagar; and 67.1 ± 4.3 SD in Black Bengal (Fig. 5B). Importantly, crossover rate across these breeds was positively correlated with both chromosome axis length (Fig. 5C and 5E) and the number of SUMO1 foci (Fig. 5D and 5F) in pachytene-stage spermatocyte nuclei. Moreover, co-staining for SUMO1 and PRR19 revealed strong positive correlations between SUMO1 foci, PRR19 foci, and axis lengths in individual spermatocyte nuclei across the five species (Fig. 5E–G and S4).

**Fig. 5.**
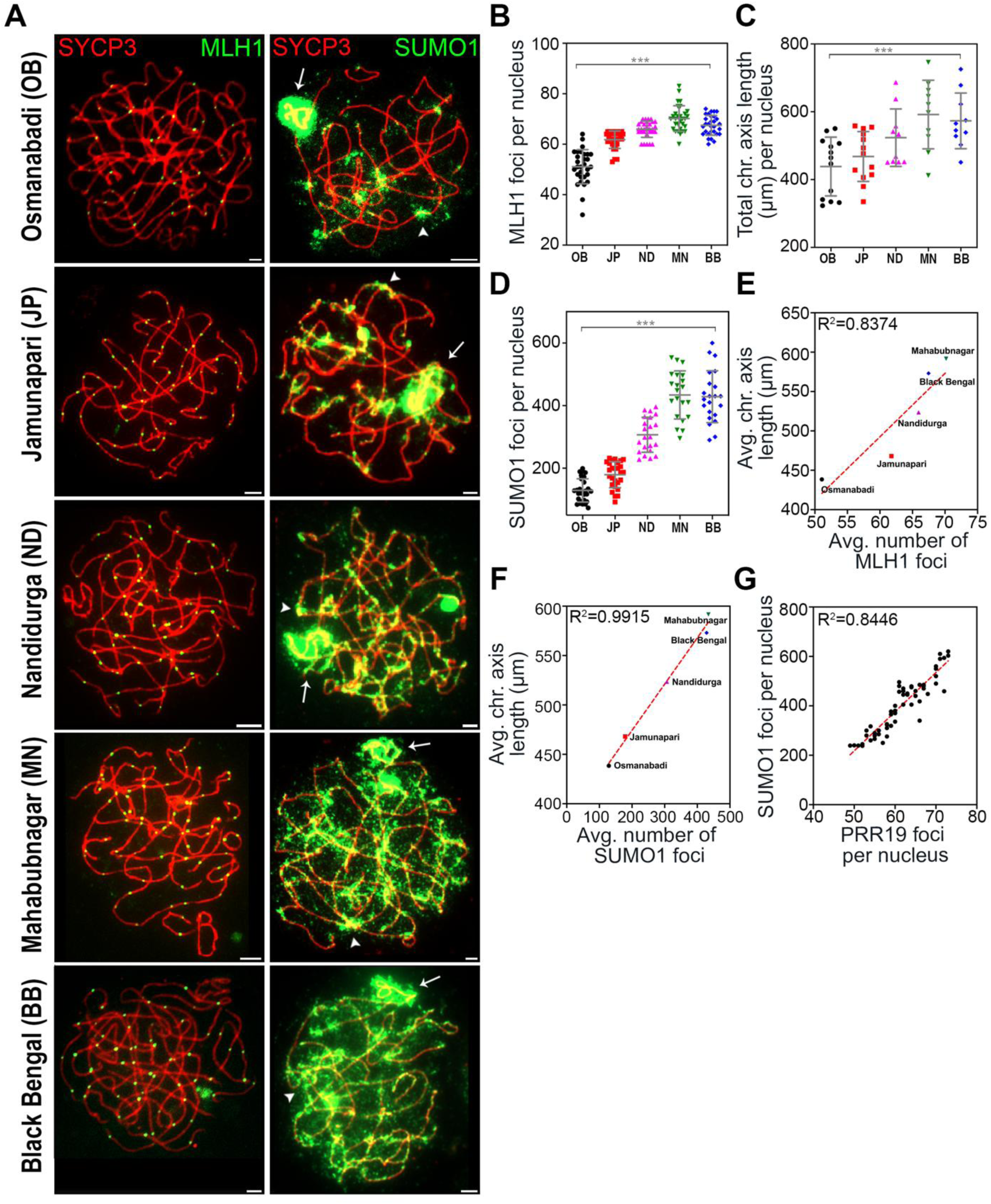
Variation in meiotic crossover and axis length across goat breeds. (**A**) Spermatocyte nuclei were immunostained for SYCP3 (red) and MLH1 (green) in respective goat breeds. (**B**) Quantification of MLH1 foci per nucleus in respective goat breeds. (**C**) Quantification of total chromosome axis length in respective goat breeds. (**D**) Quantification of SUMO1 foci per nucleus in mid pachynema nuclei. (**E**) Comparison of average chromosome axis length with average number of MLH1 foci. (**F**) Comparison of average chromosome axis length with average number of SUMO1 foci. (**G**) Correlation graph between SUMO1 and MLH1 foci per nucleus in goat spermatocytes. Error bars show mean ± s.d. (***) *P* ≤ 0.001, two-tailed Mann-Whitney. R^2^ correlation values are included in graphs along with red dashed line wherever analyzed. OB-Osmanabadi, JP-Jamunapari, ND-Nandidurga, MN-Mahabubnagar, and BB-Black Bengal. Scale bars =10μm.

### Chemical modulation of SUMO alters crossover rate in both cultured goat spermatocytes and in vivo in mouse

Collectively, our data suggest a functional relationship between SUMO levels, chromosome axis length, and crossover rate. To test this possibility, we employed short-term culture of Osmanabadi goat spermatocytes and chemical inhibitors of SUMO conjugation (the UBC9 E2-conjugase inhibitor 2-DO8) or deconjugation (the SENP1-isopeptidase inhibitor momordin)^15^ (Fig. 6A). As seen for cultured mouse spermatocytes, 2-DO8 treatment diminished chromosomal SUMO1 staining (Fig. 6B and 6C)^15^. In contrast, SUMO1 accumulated on the axes and chromatin of momordin treated spermatocytes (Fig. 6B and 6C). Analysis of MLH1 foci and chromosome axis lengths revealed opposite effects of inhibiting conjugation and deconjugation on crossover levels (Fig. 6B, 6D and 6E). In control spermatocyte nuclei (no treatment), MLH1 foci averaged 52.8 ± 4.8 SD, compared to 47.27 ± 5.5 SD in 2-DO8 treated cells, and 56 ± 6.9 SD in momordin treated cells (Fig. 6D). Total chromosome axis length in control spermatocytes averaged 380 ± 20.1 SD, compared to 340.27 ± 23.5 SD in 2-DO8 treated cells, and 421 ± 32.9 SD in momordin treated cells (Fig. 6E). These results are further evidence of a positive role for SUMO in regulating meiotic crossing over and chromosome axis length in higher-order vertebrates.

**Fig. 6.**
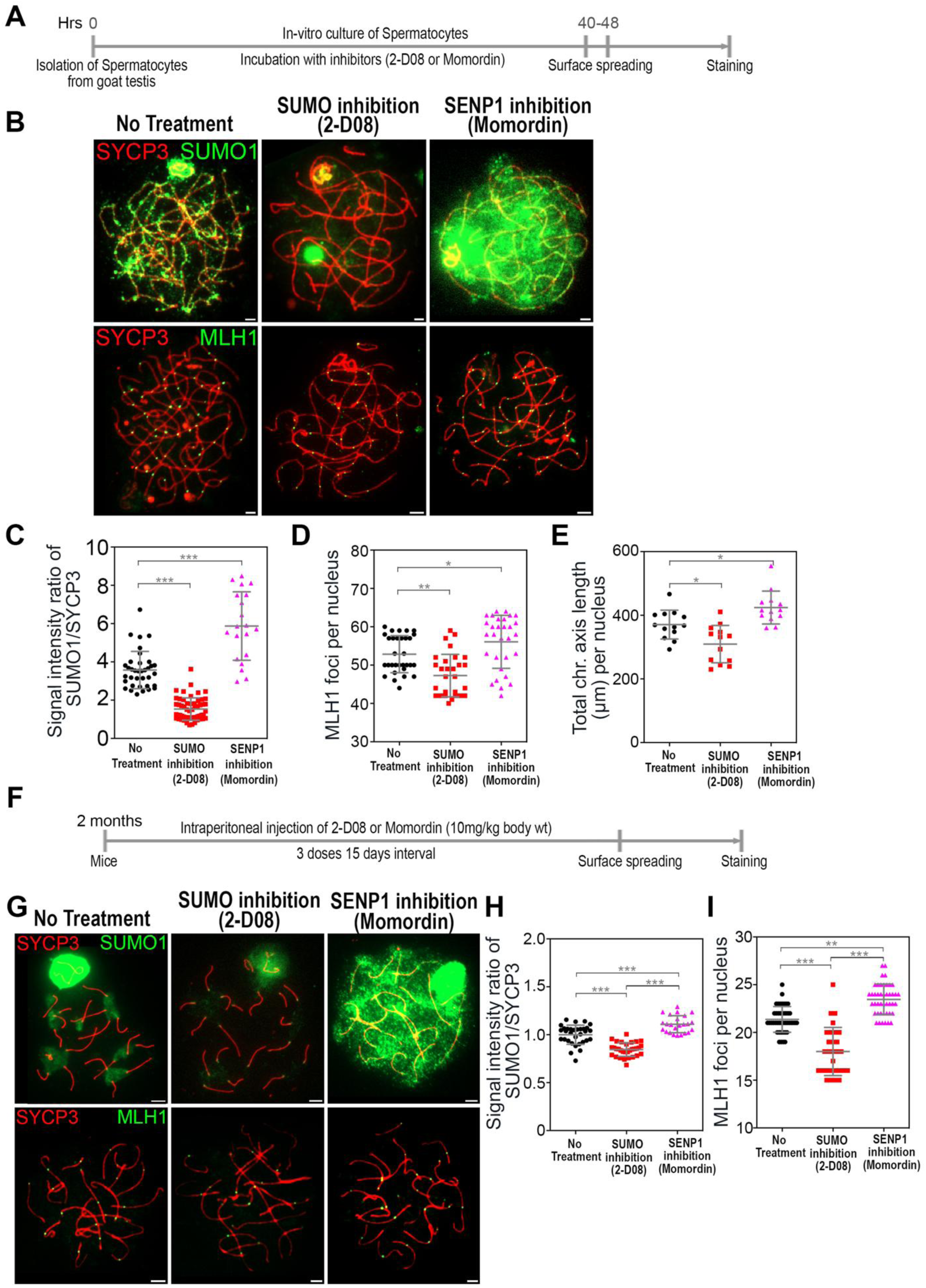
Pharmacological perturbation of SUMOylation alters meiotic recombination. **(A)** Schematic illustration of goat spermatocyte treatment with SUMO conjugation or deconjugation inhibitors. (**B**) Spermatocyte nuclei were immunostained for SYCP3 (red) and SUMO1 or MLH1 (green) in goats. (**C**) Quantification of signal intensity ratios of SUMO1/SYCP3 in respective treatments. (**D**) Quantification of MLH1 foci per nucleus in respective treatments. (**E**) Quantification of whole chromosome axis length per nucleus in respective treatments. (**F**) Schematic illustration of mice intraperitoneal injection of SUMO conjugation or deconjugation inhibitors. (**G**) Spermatocyte nuclei were immunostained for SYCP3 (red) and SUMO1 or MLH1 (green) in mice treatments. (**H**) Quantification of signal intensity ratios of SUMO1/SYCP3 in respective mice treatments. (**I**) Quantification of MLH1 foci per nucleus in respective mice treatments. Error bars show mean ± s.d. (***) *P* ≤ 0.001, (**) *P* ≤ 0.01, (*) *P* ≤ 0.1, two-tailed Mann-Whitney. Scale bars = 10μm.

These findings were further validated in mice via intraperitoneal injections of 2-DO8 or momordin. Two-month-old males were given a dose of 10 mg/kg bodyweight of either 2-DO8 or momordin every 15 days over a 45-day period (Fig. 6F). Spermatocyte chromosomes were subjected to immunofluorescence analysis to quantify SUMO levels and MLH1 foci (Fig. 6G). Similar to results obtained from cultured goat and mouse spermatocytes, 2-DO8 treatment decreased chromosomal SUMO1 staining, while momordin treatment caused SUMO1 accumulation on chromosome axes and chromatin (Fig. 6H). Untreated mouse spermatocytes averaged 22.1 ± 1.6 SD MLH1 foci, compared to 18.30 ± 2.5 SD in 2-DO8-treated cells and 24 ± 2.1 SD in momordin-treated cells (Fig. 6I).

## Discussion

Crossover rates vary significantly across species, sexes, and even individuals^26,50,51^. Such variation is thought to be shaped by evolutionary pressures, environmental factors, and genetic makeup^26,51,52^. However, the molecular mechanisms governing crossover variation remain elusive. This study provides evidence that SUMO plays a role in modulating crossover rates both between and within vertebrate species.

### Chromosome axis length as a conserved scaling parameter

Our comparative analysis shows that the positive correlation between chromosome axis length and crossover frequency, previously reported for mouse and human, can be extended across livestock vertebrate species (Fig. 1). Moreover, the quasi-fixed crossover density per unit of chromosome axis length observed in these species points to a conserved regulatory mechanism that ensures a consistent recombination rate proportional to chromosome length (Fig. 1). A likely mechanism is crossover interference, the one dimensional patterning process that reduces the occurrence of closely-spaced crossovers and results in a relatively even spacing of adjacent crossovers^53^. Our crossover density data (MLH1 foci per µm axis) imply that interference distances (MLH1 inter-focus distances) average 6.7–9.1 µm in the species studies. In this context, the chromosome axis can be viewed not merely as a scaffold but as an active regulator that spatially constrains recombination events.

However, the case of chicken highlights deviation in this model. The elevated crossover density observed in chicken is largely explained by the presence of numerous micro-chromosomes, each of which must acquire at least one crossover. This indicates that obligate crossover requirements can locally override global scaling rules, demonstrating how structural constraints and regulatory mechanisms interact to shape recombination landscapes.

### Variability in DSB/crossover ratios and DSB densities

The ratio of RAD51 foci to MLH1 foci, which reflects the number of DSBs per crossover, was relatively uniform in all species except mouse, averaging 2.8 (Fig. 2). This is equivalent to ∼35% of DSBs maturing into crossovers, which is comparable to the rate reported for human spermatocytes^27^. With ∼10 RAD51 foci per MLH1 mouse was the outlier with a much lower crossover to non-crossover ratio.

The density of RAD51 foci was similar for pig, cattle, sheep, and goat at 0.35 foci per µm of axis (Fig. 2). In contrast, chicken and mouse had significantly higher densities, at 0.82 and 1.02 foci per µm, respectively (Fig. 2). This data indicate that DSB frequency is not solely determined by axis length but can vary between species. These observations were corroborated by staining for RPA, a second marker of DSB sites marker. The relative stability of the DSB-to-crossover ratio across species indicates that only a subset of breaks is selected for crossover fate in a largely conserved manner. This shifts attention away from break formation as the dominant control point and toward later steps in recombination, where crossover designation and stabilization occur. In this light, variation in recombination rate reflects differences in how recombination intermediates are processed, rather than how frequently they are initiated.

### SUMO regulates chromosome axis length and crossover rate

One of the most notable finding is that recombination rate is not a rigid property of a species, but a plastic trait that can be tuned through changes in chromosome architecture. The strong covariation we observe between chromosome axis length, SUMO levels, and crossover frequency, both across species and within goat breeds, suggests that relatively subtle shifts in how chromosomes are organized during meiosis can produce meaningful differences in recombination output (Fig. 4 and 5). This provides a simple and elegant solution to a long-standing problem: how crossover rates can evolve without disrupting the essential requirement for at least one crossover per chromosome pair. A key feature of this model is that it decouples recombination rate from genome size and, to a large extent, from DSB formation. Instead of requiring large-scale genomic changes, recombination can be adjusted through modulation of axis length or the SUMO-dependent environment that governs recombination intermediate stability. Because the SUMO pathway targets a wide range of chromosomal proteins including cohesin, condensin, and axis components, small changes in SUMO dynamics or target specificity could alter chromosome structure in ways that scale crossover number^54,55^. In this sense, the SUMO machinery represents a broad and flexible regulatory layer capable of fine-tuning recombination.

This framework also provides a plausible explanation for the rapid evolution of recombination landscapes observed across vertebrates^56^. Changes in axis organization or SUMO regulation could shift crossover rates over relatively short evolutionary timescales, without requiring alterations to the core recombination machinery. The variation we observe among goat breeds reinforces this idea, demonstrating that such changes can arise within a single species and may contribute to differences in fertility, adaptation, or response to selection.

More broadly, our findings suggest that recombination rate is an emergent property of chromosome organization, one that can be reshaped by modifying the structural and regulatory context in which recombination occurs. By linking crossover control to chromosome architecture and SUMO-dependent regulation, this work highlights a mechanism that is both robust in maintaining meiotic fidelity and flexible in enabling evolutionary innovation.

## Materials and Methods

### Sample collection

Pig (*Sus scrofa*), cattle (*Bos indicus)*, sheep (*Ovis aries*), and goat (*Capra hircus*) testes samples were collected from the slaughterhouse. The slaughterhouse is government-licensed, and samples were collected from healthy mature animals and ready for reproduction. Three independent replicates with age-matched animals from the same breed with original breed features were used in this study. The different goat breed testes were collected from the herd in diverse geographical areas based on phenotypic characteristics in consultation with the herd’s goat herder. However, we did not get any parental history. In addition, we have confirmed the breed by phylogenetic analysis using mitochondrial hypervariable region (D-loop sequencing). Except for chicken testes, all samples were transported immediately to the lab on ice and processed by chopping seminiferous tubules for surface spreading. The leftover testes were frozen in liquid nitrogen and stored at −80 C for further use. Chicken (*Gallus gallus domesticus)* testes samples were collected in 1X PBS from a poultry farm with similar age groups and testis weights. All mice were congenic with the Balb/C background. Mice were maintained and used for experimentation according to the guidelines of the Institutional Animal Ethical Committees of the National Institute of Animal Biotechnology.

### Spreading of spermatocytes in different species

Testes were dissected from freshly killed animals, mostly from mice (2 months old), weighed, and processed for the surface spreading of spermatocyte chromosomes as described^15^. The exact process used for mice was followed to obtain chromosome spreads from goat, sheep, cattle, pig, and chicken testes with some modifications. After transporting testis samples to the lab, the samples were weighed, and the testes were chopped to get seminiferous tubules. The tubules were then fixed immediately in 1% PFA for 20-30 minutes, hypnotized by 0.75M sucrose for 5 minutes, and chopped finely to make the spermatocyte slurry, followed by spreading via hypotonic extraction buffer (HEB). After spreading, the slides were kept in a humid chamber for 3–4 hours, and then the lid was slightly opened overnight for slow drying of the slides. Following overnight drying, slides were washed once with water for 5 minutes and twice with 0.4% photoflo for 5 minutes each. Further, good-quality slides were used for immunocytochemistry.

### Goat spermatocyte culture and chemical inhibition

Short-term culture of spermatocytes was performed as described with few modifications^15^. Freshly isolated goat testes were collected from the slaughterhouse, and then chopped seminiferous tubules were transferred into a 50 ml tube containing 2 ml of TIM (Testes cell isolation medium consisting 104 mM NaCl, 45mM KCl, 1.2 mM MgSO4, 0.6 mM KH2PO4, 0.1% (w/v) glucose, 6 mM Na Lactate, 1 mM Na pyruvate). Then, 200 µl of freshly prepared collagenase (4 mg/ml of collagenase, Sigma C0130) solution was added and incubated for 50–60 min (or less, based on how well the testis tubules fall apart) at 32°C at 200 rpm with occasional inversion of the tube. After that, tubules were washed with TIM solution three times and then trypsinized with 200 µl of freshly prepared trypsin solution (1.4 mg/ml of Trypsin TRL, Worthington 3703). After digestion, 500 µl freshly prepared trypsin inhibitor solution (5 mg/ml Trypsin Inhibitor, Sigma T9003) and 50 µl DNase I solution was added. Then the spermatocytes were isolated by passing through 70, 40 µm nylon filters. Finally, the suspension was centrifuged and washed three times with TIM and once with PBS solution to remove cell debris. Pour off and remove the last bit of PBS with a pipette tip and resuspend in TCM199 media with antibiotics, also ensuring that 90% of spermatocytes are viable. Then, the cells were transferred into six wells of cell culture dishes with or without inhibitors. In culture media, Inhibitors were added at the following concentrations: 30 µM 2-D08 (Millipore; 505156; 3.7 mM stock dissolved in DMSO); 40 µM Momordin (1.3 mM stock dissolved in DMSO; PHL83281, Sigma-Aldrich). After 40–48 hrs of incubation at 37 °C with 5% CO2, cell viability was quantified using the trypan blue exclusion assay, pelleted at 900 rpm, washed with PBS, and processed for surface spreads.

### Immunofluorescent localization and co-localization of crossover proteins on synaptonemal complex axis

Immunofluorescence staining was performed as described, using the following primary antibodies with incubation overnight at room temperature: mouse anti-SYCP3 (sc-74569 Santa Cruz, 1:200 dilution), rabbit anti-SYCP3 (NB300-232 Gene biological solutions, 1:200 dilution), goat-anti-SYCP3 (AF3750 R&D Systems, 1:200), mouse monoclonal anti -MLH1 (3515S CST, 1:30) mouse anti-Rad51 (MA5-14419, 1:100), rabbit monoclonal anti-RPA32 (ab76420 Abcam, 1:100), mouse anti-SUMO1 (DSHB-S1-1456, 1:100), goat anti-RNF212 (gift from Neil Hunter), guina pig anti-PRR19 (kind gift from Dr. Attila Toth). After incubation, slides were washed thrice with TBST for 5 min each, blocked with ADB (Antibody dilution buffer) for 15 min, and then stained with secondary antibodies. Slides were subsequently incubated with the following goat secondary antibodies for 1hr at 37°C: Donkey anti-goat 555 (A-21432, 1:2000), Donkey anti-goat 350 (A-21081, 1:1000), Goat anti-mouse 488 (A32723, 1:1500), Goat anti-mouse 555 (A32727, 1:2000), Goat anti-rabbit 555 (A32732, 1:2000), Goat anti-rabbit 488 (A32731, 1:1500), Goat anti-guinea pig 594 (A11076, 1:1500). Coverslips were mounted with Prolong Diamond antifade reagent (Molecular Probes). For SUMO1 staining, TIM spreads were used as described^57^. Spermatocyte spread slides were incubated with pepsin buffer (0.01N HCl) for 2 min, then treated with 5 µg/ml pepsin in 0.01N HCl for 3 min at room temperature, followed by 5 min wash with 1X TBST. Slides were then incubated in DNase buffer for 2 min, treated with 1:100 (1 mg/ml) DNase for 20 min at 37°C. After DNase treatment, slides were washed with 1X TBST for 5 min, blocked with ADB for twice 15 min each, and then proceeded for primary antibody staining.

### Imaging and counting of imaged data

Immunolabeled chromosome-spread nuclei were imaged using a Zeiss Axio Observer 7 microscope with a 100× Plan Apochromat 1.4 NA objective and HXP 120V X-Cite 120 metal halide light source. Images were captured by an Axiocam 503 CCD camera, processed using Zeiss software, and analyzed with Zen 2.6 blue software.

Late pachytene spermatocytes were selected for MLH1 foci quantification, with staging based on sex chromosome morphology. The mean number of MLH1 foci per nucleus was calculated and plotted against genome size (Table 1; Fig. 1D) and chromosome number (Table 1; Fig. 1F). In addition, MLH1 foci counts were analyzed in relation to chromosome arm number (Table 1), which was determined based on centromere position: metacentric, submetacentric, subtelocentric and acrocentric chromosomes were each considered to have two arms, whereas telocentric chromosomes were scored as having a single arm. Chromosome axis lengths were measured manually using ZEN Blue software (Zeiss). The synaptonemal complex was traced using the distance spline tool based on SYCP3 immunostaining, and the lengths of all chromosome axes within a nucleus were summed to obtain the total synaptonemal complex length per nucleus.

Two observers performed all cytological analyses; the second observer was blind to which group was being analyzed. SYCP3-staining nuclei were staged using standard criteria. For quantification of all the markers, only foci closely associated with (“touching”) the SYCP3-staining chromosome axes were counted.

### Pharmacological treatments in mice

Two-month-old male mice were administered intraperitoneal injections of either 2-DO8 or momordin at a dose of 10 mg/kg body weight, every 15 days over 45 days. Following the treatment period, the mice were sacrificed, and the testes were collected for spermatocyte spreading using the HEB and TIM procedures. The spread slides were used for immunostaining as described above.

### Data acquisition and phylogeny approach

Phylogenetic analysis of the meiosis-specific gene sequences was done using MEGA X^58^. Initially, the sequence alignments were done using MUSCLE^59^. The aligned sequences were analyzed for the best nucleotide substitution model based on Bayesian information criterion scores using the JModel Test software v2.1.7, and the cladogram was drawn by BEAST 2.0^60,61^. The tree was constructed by the Neighbor-joining method with the best model obtained, using 1000 bootstrap replicates.

## ACKNOWLEDGMENTS

We thank Attila Tóth for kindly donating antibodies. We are grateful to the NIAB Core Microscopy Facility for imaging support. L.K.S. was supported by a DBT Junior Research Fellowship (JRF). R.B. was supported by a UGC Junior Research Fellowship (JRF), and A.M. was supported by a CSIR Junior Research Fellowship (JRF). This work was supported by NIAB core funds and the DBT Ramalingaswami Re-entry Fellowship (BT/RLF/Re-entry/21/2016), as well as DBT grant BT/PR46563/AAQ/1/868/2022 awarded to H.B.D.P.R.

## AUTHOR CONTRIBUTIONS

S.L.K. and H.B.D.P.R. conceived the study and designed the experiments. S.L.K., R.B., A.M., A.K., A.K., G.R.K., and H.B.D.P.R. performed the experiments and analyzed the data. S.L.K., N.H., and H.B.D.P.R. wrote the manuscript with inputs and edits from all authors.

## DATA AVAILABILITY STATEMENT

The data underlying this article are available in the article and in its online supplementary material.

## DECLARATION OF INTERESTS

The authors declare no competing interests.

None of the material reported in this manuscript has been published or made available on the Internet, nor is it under consideration elsewhere. Animals used for experimentation according to the guidelines of the Institutional Animal Ethics Committees of the National Institute of Animal Biotechnology, Hyderabad, India.

## Supplementary figure legends

**Fig. S1.**
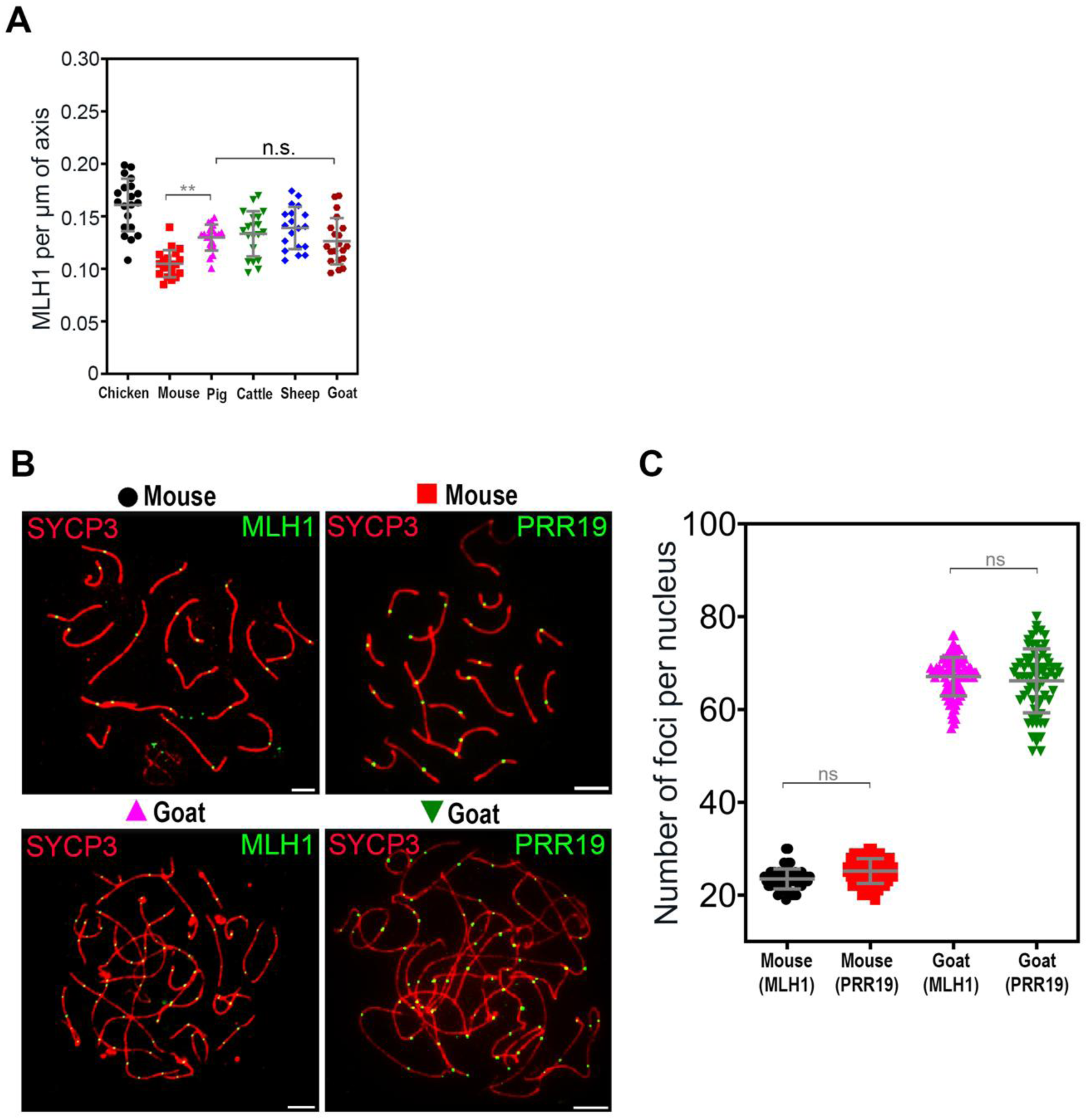
Differential regulation of micro-chromosome crossover in chicken. (**A**) Quantification of MLH1 per micron of chromosome axis, excluding chicken micro chromosomes. (**B**) Spermatocyte nuclei of mouse and goat species immunostained for axis marker SYCP3 (red) and the crossover marker MLH1 (green) or PRR19 (green). (**C**) Quantification of immunostaining MLH1 or PRR19 foci in both mouse and goat species. Error bars show mean ± s.d. (**) *P* ≤ 0.01, (n.s.) *P* ≥ 0.1, two-tailed Mann-Whitney.

**Fig. S2.**
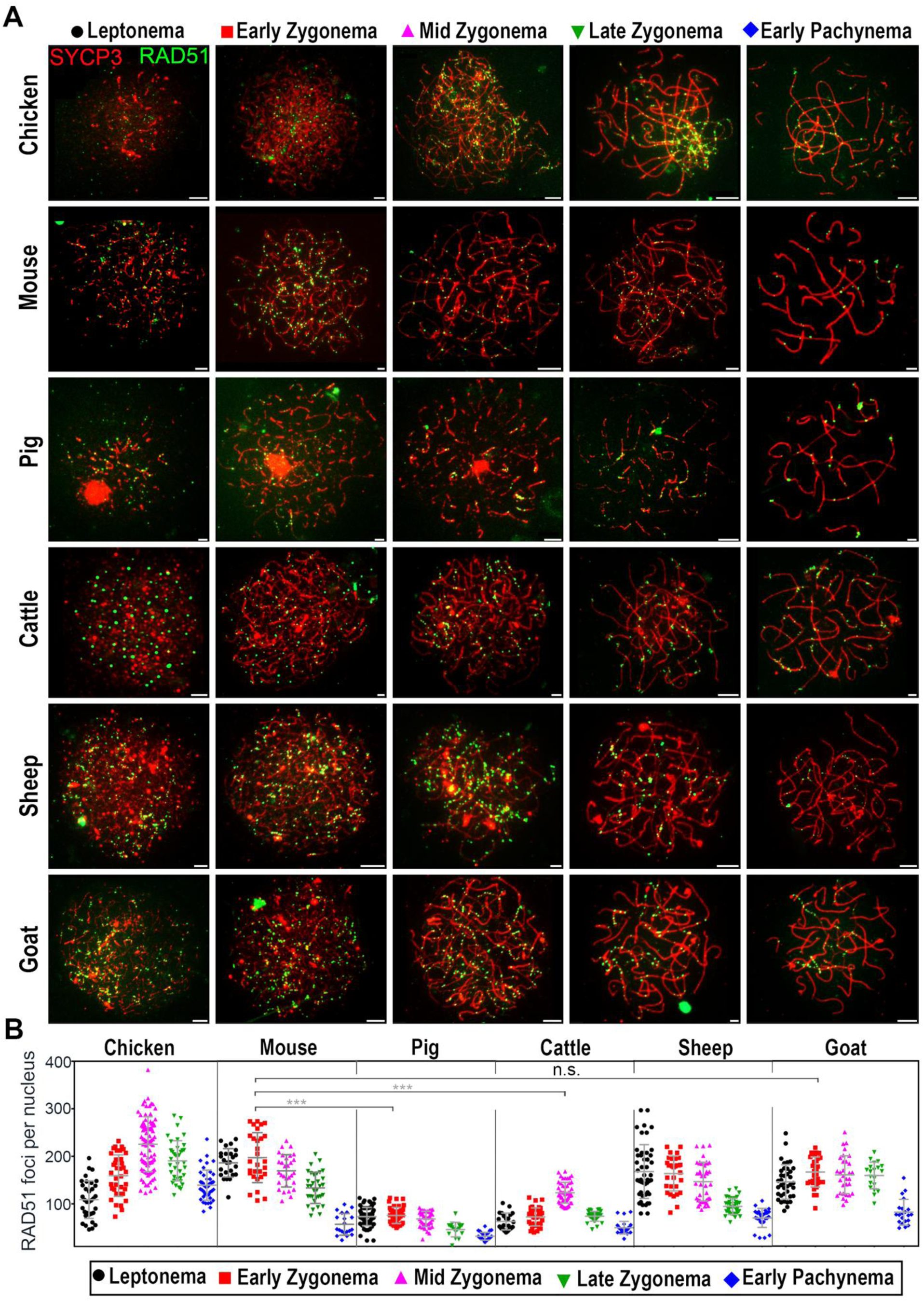
RAD51 dynamics in vertebrates: **(A)** Spermatocyte nuclei from the indicated species immunostained for SYCP3 (red) and RAD51 (green) in respective stages. (**B**) Quantification of immunostaining foci in respective stages. Error bars show mean ± s.d. (***) *P* ≤ 0.001, (n.s.) *P* ≥ 0.1, two-tailed Mann-Whitney. Scale bars = 10μm.

**Fig. S3.**
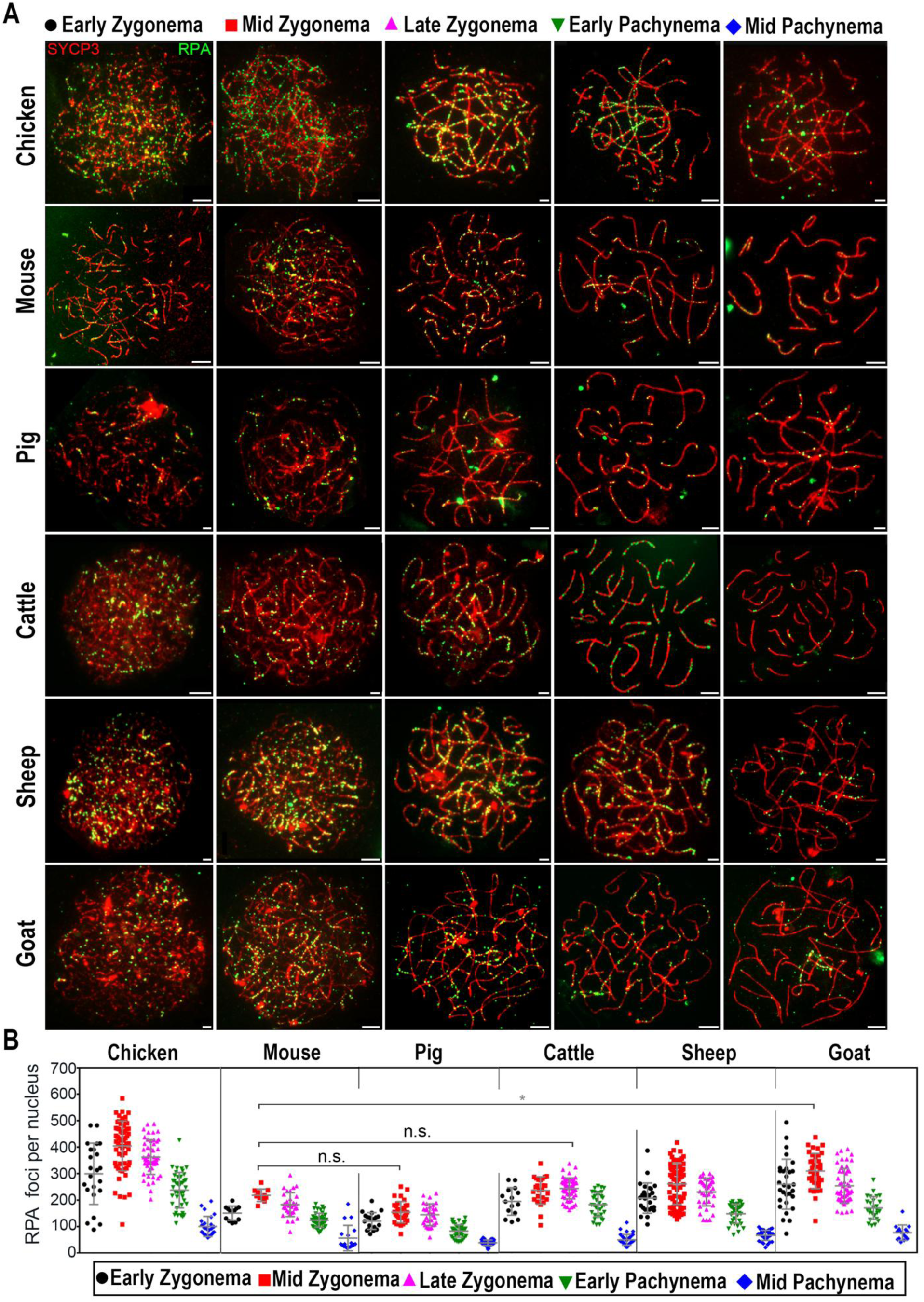
RPA dynamics in vertebrates: **(A)** Spermatocyte nuclei from the indicated species immunostained for SYCP3 (red) and RPA (green) in respective stages. (**B**) Quantification of immunostaining foci in respective stages. Error bars show mean ± s.d. (*) *P* ≤ 0.1, (n.s.) *P* ≥ 0.1, two-tailed Mann-Whitney. Scale bars = 10μm.

**Fig. S4.**
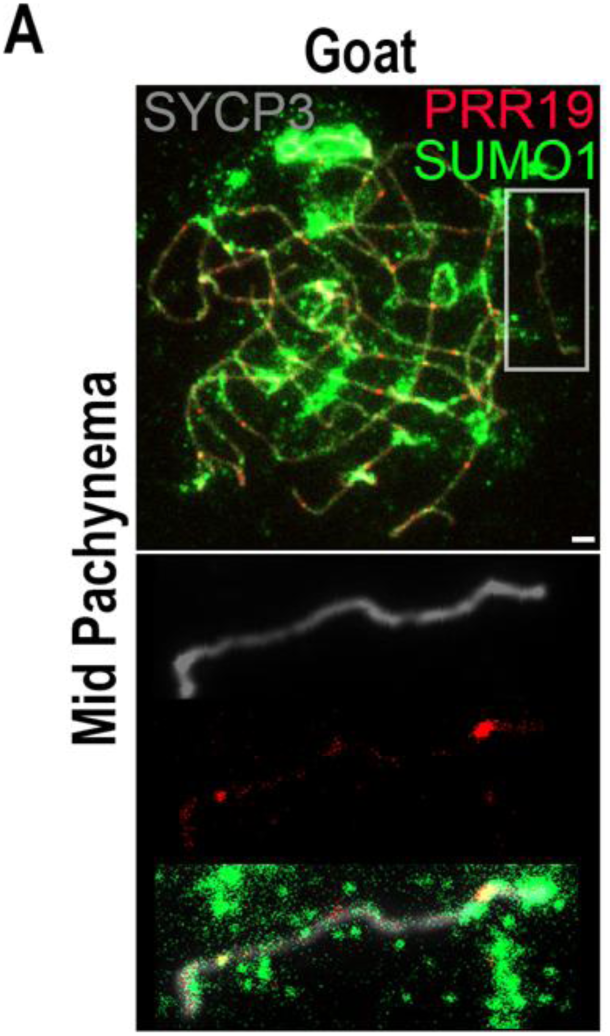
SUMO and PRR19 colocalization in goat: Spermatocyte nuclei from goat immunostained for SYCP3 (grey), SUMO1 (green) and PRR19 (red) in mid-pachytene stage. Zoom in on the chromosome shown below. Scale bars = 10μm.

